# Using GPT-4 to write a scientific review article: a pilot evaluation study

**DOI:** 10.1101/2024.04.13.589376

**Authors:** Zhiping Paul Wang, Priyanka Bhandary, Yizhou Wang, Jason H. Moore

**Affiliations:** Department of Computational Biomedicine, Cedars Sinai Medical Center, 700 N. San Vicente Blvd., Pacific Design Center, Suite G-541, West Hollywood, 90069, CA, USA

## Abstract

GPT-4, as the most advanced version of OpenAI’s large language models, has attracted widespread attention, rapidly becoming an indispensable AI tool across various areas. This includes its exploration by scientists for diverse applications. Our study focused on assessing GPT-4’s capabilities in generating text, tables, and diagrams for biomedical review papers. We also assessed the consistency in text generation by GPT-4, along with potential plagiarism issues when employing this model for the composition of scientific review papers. Based on the results, we suggest the development of enhanced functionalities in ChatGPT, aiming to meet the needs of the scientific community more effectively. This includes enhancements in uploaded document processing for reference materials, a deeper grasp of intricate biomedical concepts, more precise and efficient information distillation for table generation, and a further refined model specifically tailored for scientific diagram creation.

## Introduction

A comprehensive review of a research field can significantly aid researchers in quickly grasping the nuances of a specific domain, leading to well-informed research strategies, efficient resource utilization, and enhanced productivity. However, the process of writing such reviews is intricate, involving multiple time-intensive steps. These include the collection of relevant papers and materials, the distillation of key points from potentially hundreds or even thousands of sources into a cohesive overview, the synthesis of this information into a meaningful and impactful knowledge framework, and the illumination of potential future research directions within the domain. Given the breadth and depth of biomedical research—one of the most expansive and dynamic fields—crafting a literature review in this area can be particularly challenging and time-consuming, often requiring months of dedicated effort from domain experts to sift through the extensive body of work and produce a valuable review paper [1,2].

The swift progress in Natural Language Processing (NLP) technology, particularly with the rise of Generative Pre-trained Transformers (GPT) and other Large Language Models (LLMs), has equipped researchers with a potent tool for swiftly processing extensive literature. A recent survey indicates that ChatGPT has become an asset for researchers across various fields [3]. For instance, a PubMed search for “ChatGPT” yielded over 1,400 articles with ChatGPT in their titles as of November 30th, 2023, marking a significant uptake just one year after ChatGPT’s introduction.

The exploration of NLP technology’s capability to synthesize scientific publications into comprehensive reviews is ongoing. The interest in ChatGPT’s application across scientific domains is evident. Studies have evaluated ChatGPT’s potential in clinical and academic writing [3-10], and discussions are underway about its use as a scientific review article generator [11,12,13]. However, many of these studies predate the release of the more advanced GPT-4, which may render their findings outdated. In addition, there is no study specifically evaluating ChatGPT (GPT-4) for writing biomedical review papers.

As the applications of ChatGPT are explored, the scientific community is also examining the evolving role of AI in research. Unlike any tool previously utilized in the history of science, ChatGPT has been accorded a role akin to that of a scientist, even being credited as an author in scholarly articles [14]. This development has sparked ethical debates. While thorough evaluations of the quality of AI-generated scientific review articles are yet to be conducted, some AI tools, such as Scopus AI [15], are already being employed to summarize and synthesize knowledge from scientific literature databases. However, these tools often come with disclaimers cautioning users about the possibility of AI generating erroneous or offensive content. Concurrently, as ChatGPT’s potential contributions to science are probed, concerns about the possible detrimental effects of ChatGPT and other AI tools on scientific integrity have been raised [16]. These considerations highlight the necessity for more comprehensive evaluations of ChatGPT from various perspectives.

In this study, we evaluated the capabilities of ChatGPT with GPT-4 in writing biomedical review papers, using two cancer research papers as benchmarks. The first paper [17] served as a test for ChatGPT’s ability to generate main points and summarize text, while the second paper [18] tested its capacity for creating tables and graphs. We simulated the steps a scientist would take in writing a cancer research review and assessed GPT-4’s performance at each stage. Our findings are presented across four dimensions: 1) the ability to summarize insights from reference papers on specific topics; 2) the semantic similarity of GPT-4 generated text to benchmark texts; 3) the projection of future research directions based on current publications; and 4) the synthesis of context in the form of tables and graphs. We conclude with a discussion of our overall experience and the insights gained from this study.

## Method

### Review text content generation by ChatGPT

The design of this study aims to replicate the process a scientist undergoes when composing a biomedical review paper. This involves the meticulous collection, examination, and organization of pertinent references, followed by the articulation of key topics of interest into a structured format of sections, subsections, and main points. The scientist then synthesizes information from the relevant references to develop a comprehensive narrative. A primary objective of this study is to assess ChatGPT’s proficiency in distilling insights from references into coherent text. To this end, a review paper on sex differences in cancer [17] was chosen as a benchmark, referred to as BRP1 (Benchmark

Review Paper 1). Using BRP1 for comparison, ChatGPT’s content generation was evaluated across three dimensions: 1) summarization of main points; 2) generation of review content for each main point; and 3)synthesis of information from references to project future research directions.

#### Main point summarization

The effectiveness of GPT-4 in summarizing information was tested by providing it with the 113 reference articles from BRP1 to generate a list of potential sections for a review paper. The generated sections

were then compared with BRP1’s actual section titles for coverage evaluation (Figure 1(A)). Additionally, GPT-4 was tasked with creating possible subsections using the BRP1 section titles and reference articles, which were compared with the actual subsection titles in BRP1.

**Figure 1.**
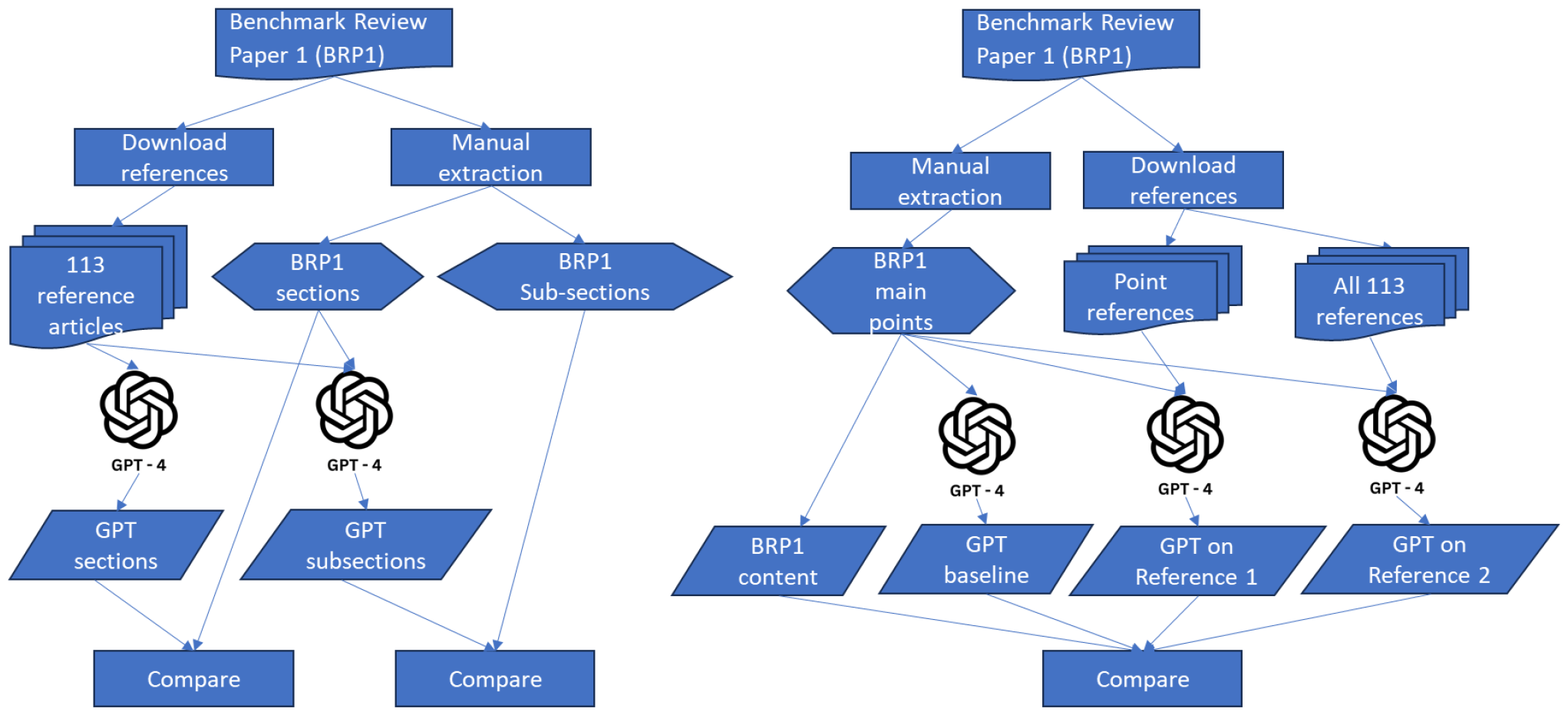
(A) GPT-4 summarizes sections and subsections; (B) GPT-4 generated review content evaluation

#### Review content generation

The review content generation test involved comparing GPT-4’s ability to summarize a given point with the actual text content from BRP1 (Figure 1(B)). BRP1 comprises three sections with seven subsections, presenting a total of eight main points. The corresponding text content for each point was manually extracted from BRP1. Three strategies were employed for GPT-4 to generate detailed elaborations for these main points: 1) providing a point only in a prompt for baseline content generation; 2) feeding all references used by BRP1 to GPT-4 for reference-based content generation; 3) using only the references corresponding to a main point, i.e., articles being referred in a subsection of BRP1, for content generation to make a main point. The semantic similarity of the text content generated by these strategies was then compared with the manually extracted content from BRP1.

#### Projections on future research

The section on “outstanding questions” in the Concluding Remarks of BRP1 serves a dual purpose: it summarizes conclusions and sets a trajectory for future research into sex differences in cancer. This is a common feature in biomedical review papers, where a forward-looking analysis is synthesized from the main discussions within the paper. The pivotal inquiry is whether ChatGPT, without further refinement, can emulate this forward projection using all referenced articles. The relevance of such a projection is contingent upon its alignment with the main points and references of the review. Moreover, it raises the question of whether the baseline GPT-4 LLM would perform comparably.

To address these queries, all references from BRP1 were inputted into GPT-4 to generate a section akin to Concluding Remarks, encompassing a description of sex differences in cancer, future work, and potential research trajectories. Additionally, three distinct strategies were employed to assess GPT-4’s ability to formulate specific “outstanding questions,” thereby evaluating ChatGPT’s predictive capabilities for future research. These strategies involved uploading all BRP1 reference articles to GPT-4 for projection: 1) without any contextual information; 2) with the inclusion of BRP1’s main points; 3) with a brief description of broad areas of interest. The outputs from these strategies, along with the base model’s output—GPT-4 without reference articles—were juxtaposed with BRP1’s original “outstanding questions” for comparison.

### Data process

#### ChatGPT query

In initiating this study, we utilized the ChatGPT web application (https://chat.openai.com/). However, we encountered several limitations that impeded our progress:

1. A cap of ten file uploads, which restricts the analysis of content synthesized from over ten articles.
2. A file size limit of 50MB, hindering the consolidation of multiple articles into a single file to circumvent the upload constraint.
3. Inconsistencies in text file interpretation when converted from PDF format, rendering the conversion of large PDFs to smaller text files ineffective.
4. Anomalies in file scanning, where ChatGPT would occasionally process only one of several uploaded files, despite instructions to utilize all provided files.

Due to these constraints, we transitioned to using GPT-4 API calls for all tests involving document processing. The GPT-4 API accommodates up to twenty file uploads simultaneously, efficiently processes text files converted from PDFs, and demonstrates reliable file scanning for multiple documents. The Python code, ChatGPT prompts, and outputs pertinent to this study are available in the supplementary materials.

#### Text similarity comparison

To assess text content similarity, we employed a transformer network-based pre-trained model [19] to calculate the semantic similarity between the original text in BRP1 and the text generated by GPT-4. We utilized the *util*.*pytorch_cos_sim* function from the *sentence_transformers* package to compute the cosine similarity of semantic content. Additionally, we conducted a manual validation to categorize the similarity between the GPT-4 generated content and the original BRP1 content into three distinct levels: semantically very similar (Y), partially similar (P), and not similar (N).

### Reproducibility and Plagiarism evaluation

The inherent randomness in ChatGPT’s output, attributable to the probabilistic nature of large language models (LLMs), necessitates the validation of reproducibility for results derived from ChatGPT outputs. To obtain relatively consistent responses from ChatGPT, it is advantageous to provide detailed context within the prompt, thereby guiding the model towards the desired response. Consequently, we replicated two review content generation tests, as depicted in Figure 1(B)—one based on point references and the other on the GPT-4 base model—one week apart using identical reference articles and prompts via API calls to GPT-4. The first test aimed to evaluate the consistency of file-based content generation by GPT-4, while the second assessed the base model. We compared the outputs from the subsequent run with those from the initial run to determine the reproducibility of the text content generated by ChatGPT.

Prior to considering the utilization of ChatGPT for generating content suitable for publication in a review paper, it is critical to address potential plagiarism concerns. The pivotal question is whether text produced by GPT-4 would be flagged as plagiarized by anti-plagiarism software. In this study, GPT-4 generated a substantial volume of text, particularly for the text content comparison test (Figure 1(B)).

We subjected both the base model-generated review content and the reference-based GPT-4 review content to scrutiny using iThenticate to ascertain the presence of plagiarism.

### Table and figure generation by ChatGPT

Review papers often distill the content from references into tables and further synthesize this information into figures. In this study, we evaluated ChatGPT’s proficiency in generating content in tabular and diagrammatic formats, using benchmark review paper 2 (BRP2) [18] as a reference, as illustrated in Figure 2. The authors of BRP2 developed the seminal Cancer-Immunity Cycle concept, encapsulated in a cycle diagram, which has since become a structural foundational for research in cancer immunotherapy.

**Figure 2.**
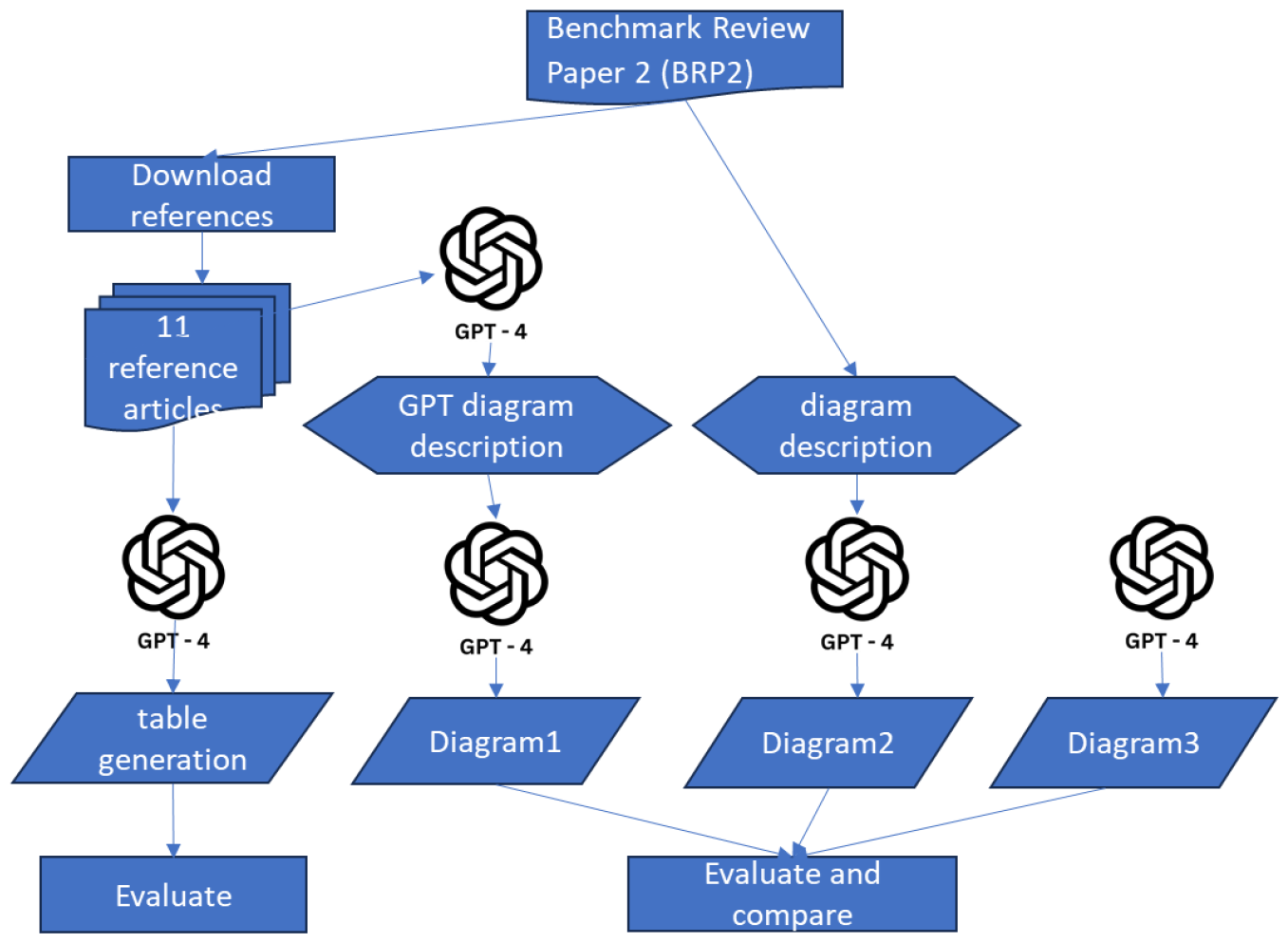
GPT-4 table generation and figure creation

#### Table content generation

Analogous to the file scan anomaly, ChatGPT may disproportionately prioritize one task over others when presented with multiple tasks simultaneously. To mitigate this in the table generation test, we adopted a divide-and-conquer approach, submitting separate GPT-4 prompts to generate content for each column of the table. This strategy facilitated the straightforward assembly of the individual outputs into a comprehensive table, either through GPT-4 or manual compilation.

In BRP2, eleven reference articles were utilized to construct a table (specifically, Table 1 of BRP2) that categorized positive and negative regulators at each stage of the Cancer-Immunity Cycle. These articles were compiled and inputted into ChatGPT, prompting GPT-4 to summarize information for corresponding table columns: *Steps, Stimulators, Inhibitors, Other Considerations*, and *Example References*. The content for each column was generated through separate GPT-4 API calls and subsequently compared manually with the content in the original BRP2 table.

**Table 1.** evaluation of GPT-4 generated content by comparing with the corresponding text from the original review paper (BRP1).

| Point index | Similarity score 1<br>(use corresponding references only) | Similarity score 1<br>manual check<br>(Y/P/N) | Similarity score 2<br>(use all references from the review paper) | Similarity score 2<br>manual check<br>(Y/P/N) | Similarity score 3<br>(use GPT-4 base model only) | Similarity score 3<br>manual check<br>(Y/P/N) |
| --- | --- | --- | --- | --- | --- | --- |
| 1 | 0.707 | P | 0.632 | P | 0.593 | P |
| 2 | 0.782 | Y | 0.602 | P | 0.763 | Y |
| 3 | 0.757 | Y | 0.826 | Y | 0.823 | Y |
| 4 | 0.843 | Y | 0.811 | Y | 0.799 | Y |
| 5 | 0.740 | Y | 0.705 | Y | 0.801 | Y |
| 6 | 0.620 | P | 0.509 | P | 0.671 | P |
| 7 | 0.762 | Y | 0.739 | Y | 0.790 | Y |
| 8 | 0.771 | Y | 0.770 | Y | 0.801 | Y |
| Summary | 0.748 | 6Y2P | 0.699 | 5Y3P | 0.755 | 6Y2P |

**Table 2.** GPT-4 projection performance.

| GPT-4 query type | Similarity score 1<br>(all references) | Similarity score 1<br>manual check<br>(Y/P/N) | Similarity score 2<br>(GPT-4 base<br>model) | Similarity score 2<br>manual check<br>(Y/P/N) |
| --- | --- | --- | --- | --- |
| content summary | 0.772 | Y | 0.783 | Y |
| projection without context | 0.450 | P | 0.717 | Y |
| projection with given main points | 0.714 | P | 0.675 | P |
| projection with interested direction | 0.788 | Y | 0.760 | Y |

#### Diagram creation

ChatGPT is primarily designed for text handling, yet its capabilities in graph generation are increasingly being explored [20]. DALL-E, the model utilized by ChatGPT for diagram creation, has been trained on a diverse array of images, encompassing various subjects, styles, contexts, and including scientific and technical imagery. To direct ChatGPT towards producing a diagram that closely aligns with the intended visualization, a precise and succinct description of the diagram is essential. Like the approach for table generation, multiple prompts may be required to facilitate incremental revisions in the drawing process.

In this evaluation, we implemented three distinct strategies for diagram generation, as demonstrated in Figure 2. Initially, the 11 reference articles used for table generation were also employed by GPT-4 to generate a description for the cancer immunity cycle, followed by the creation of a diagrammatic representation of the cycle by GPT-4. This approach not only tested the information synthesis capability of GPT-4 but also its diagram drawing proficiency. Secondly, we extracted the paragraph under the section titled ‘The Cancer-Immunity Cycle’ from BRP2 to serve as the diagram description. Terms indicative of a cyclical structure, such as ‘cycle’ and ‘step 1 again,’ were omitted from the description prior to its use as a prompt for diagram drawing. This tested GPT-4’s ability to synthesize the provided information into an innovative cyclical structure for cancer immunotherapy. Lastly, the GPT-4 base model was tasked with generating a cancer immunity mechanism and its diagrammatic representation without any given context. The diagrams produced through these three strategies were scrutinized and compared with the original cancer immunity cycle figure in BRP2 to assess the scientific diagram drawing capabilities of GPT-4

## Results and Discussions

### Review content generation

#### Main point summary

As depicted in Figure 1A, GPT-4 generated nine potential sections for a proposed paper entitled ‘The Spectrum of Sex Differences in Cancer,’ utilizing the 113 reference articles uploaded, which encompassed all three sections in BRP1. Upon request to generate possible subsections using BRP1 section titles and references, GPT-4 produced four subsections for each section, totaling twelve subsections that encompassed all seven subsections in BRP1. Detailed information regarding GPT-4 prompts, outputs, and comparisons with BRP1 section and subsection titles is provided in the supplementary materials.

The results suggest that ChatGPT can effectively summarize the key points from a comprehensive list of documents, which is particularly beneficial when composing a review paper that references hundreds of articles. With ChatGPT’s assistance, authors can swiftly summarize a list of main topics for further refinement, organization, and editing. Once the topics are finalized, GPT-4 can easily summarize different aspects for each topic, aiding authors in organizing the subsections. This indicates a novel approach to review paper composition that could be more efficient and productive than traditional methods. It represents a collaborative effort between ChatGPT and the review writer, with ChatGPT sorting and summarizing articles, and the author conducting high-level and creative analysis and editing.

During this evaluation, one limitation of GPT-4 was identified: its inability to provide an accurate list of articles referenced for point generation. This presents a challenge in developing an automated pipeline that enables both information summarization and file classification.

#### Review content generation

The evaluation results for GPT-4’s review content generation are presented in Table 1 (refer to Figure 1B). When generating review content using corresponding references as in BRP1, GPT-4 achieved an average similarity score of 0.748 with the original content in BRP1 across all main points. Manual similarity validation confirmed that GPT-4 generated content that was semantically similar for all 8 points, with 6 points matching very well (Y) and 2 points matching partially (P). When utilizing all reference articles for GPT-4 to generate review content for a point, the mean similarity score was slightly lower at 0.699, with a manual validation result of 5Y3P. The results from the GPT-4 based model were comparable to the corresponding reference-based results, with a mean similarity score of 0.755 and a 6Y2P manual validation outcome.

As the GPT-4 base model has been trained on an extensive corpus of scientific literature, including journals and articles that explore sex differences in cancer, it is plausible for it to generate text content similar to the original review paper, even for a defined point without any contextual input. The performance when using corresponding references is notably better than when using all references, suggesting that GPT-4 processes information more effectively with relevant and less noisy input.

The similarity score represents only the level of semantic similarity between the GPT-4 output and the original review paper text. It should not be construed as a measure of the quality of the text content generated by GPT-4. While it is relatively straightforward to assess the relevance of content for a point, gauging comprehensiveness is nearly impossible without a gold standard. However, scientific review papers are often required in research areas where such standards do not yet exist. Consequently, this review content similarity test merely indicates whether GPT-4 can produce text content that is semantically akin to that of a human scholar. Based on the results presented in Table 1, GPT-4 has demonstrated adequate capability in this regard.

#### Projection

In this evaluation, GPT-4 initially synthesized content analogous to the *Concluding Remarks* section of BRP1 by utilizing all reference articles, further assessing its capability to integrate information into coherent conclusions. Subsequently, GPT-4 projected future research directions using three distinct methodologies. When provided with all references, GPT-4 demonstrated improved performance in content generation, suggesting that more relevant information serves as a better guide for the model. In contrast, the performance of the GPT-4 base model remained comparably stable, regardless of additional contextual cues. Manual verification confirmed GPT-4’s ability to synthesize information from the provided documents and to make reasonably accurate predictions about future research trajectories.

### Reproducibility

The comparative analysis of GPT-4 outputs from different runs is presented in Table 3. Based on previous similarity assessments, a similarity score of 0.7 is generally indicative of a strong semantic correlation in the context of this review paper. In this instance, GPT-4 outputs using corresponding references exhibited an average similarity score of 0.8 between two runs, while the base model scored

**Table 3.** GPT-4 text content reproducibility evaluation.

| Point index | Similarity score 1 (corresponding references only) | Similarity score 1 manual check (Y/P/N) | Similarity score 2 (GPT-4 base model) | Similarity score 2 manual check (Y/P/N) |
| --- | --- | --- | --- | --- |
| 1 | 0.842 | Y | 0.913 | Y |
| 2 | 0.793 | Y | 0.912 | Y |
| 3 | 0.810 | Y | 0.902 | Y |
| 4 | 0.820 | Y | 0.948 | Y |
| 5 | 0.683 | Y | 0.900 | Y |
| 6 | 0.794 | Y | 0.862 | Y |
| 7 | 0.856 | Y | 0.914 | Y |
| 8 | 0.860 | Y | 0.912 | Y |
| Summary | 0.807 | 8Y | 0.908 | Y |

0.9. A manual review confirmed that both outputs expressed the same semantic meaning at different times. Consequently, it can be concluded that GPT-4 consistently generates uniform text responses when provided with identical prompts and reference materials.

An intriguing observation is that the GPT-4 base model appears to be more stable than when utilizing uploaded documents. This may suggest limitations in GPT-4’s ability to process external documents, particularly those that are unstructured or highly specialized in scientific content. This limitation aligns with our previous observation regarding GPT-4’s deficiency in cataloging citations within its content summaries.

### Plagiarism check

The plagiarism assessment conducted via iThenticate (https://www.ithenticate.com/) yielded a percentage score of 34% for reference-based GPT-4 content generation and 10% for the base model. Of these percentages, only 2% and 3%, respectively, were attributed to matches with the original review paper (BRP1), predominantly due to title similarities, as we maintained the same section and subsection titles. A score of 34% is typically indicative of significant plagiarism concerns, whereas 10% is considered minimal. These results demonstrate the GPT-4 base model’s capacity to expound upon designated points in a novel manner, minimally influenced by the original paper. However, the reference-based content generation raised concerns due to a couple of instances of ‘copy-paste’ style matches from two paragraphs in BRP1 references [21,22], which contributed to the elevated 34% score. In summary, while the overall content generated by ChatGPT appears to be novel, the occurrence of sporadic close matches warrants scrutiny.

This finding aligns with the theoretical low risk of direct plagiarism by ChatGPT, as AI-generated text responses are based on learned patterns and information, rather than direct ‘copy-paste’ from specific sources. Nonetheless, the potential for plagiarism and related academic integrity issues are of serious concern in academia. Researchers have been exploring appropriate methods to disclose ChatGPT’s contributions in publications and strategies to detect AI-generated content [23,24,25]

### Table content generation

Table construction in scientific publications often necessitates a more succinct representation of relationships and key terms compared to text content summarization and synthesis. This requires ChatGPT to extract information with greater precision. For the five columns of information compiled by GPT-4 for Table 1 in BRP2, the *Steps* column is akin to summarizing section and subsection titles in BRP1. ‘*Stimulators’* and ‘*Inhibitors’* involve listing immune regulation factors, demanding more concise and precise information extraction. ‘*Other Considerations*’ encompasses additional relevant information, while ‘*Example References*’ lists citations.

For the *Steps* column, GPT-4 partially succeeded but struggled to accurately summarize information into numbered steps. For the remaining columns, GPT-4 was unable to extract the corresponding information accurately. Extracting concise and precise information from uploaded documents for specific scientific categories remains a significant challenge for GPT-4, which also lacks the ability to provide reference citations, as observed in previous tests. All results, including GPT prompts, outputs, and evaluations, are detailed in the supplementary materials.

In summary, GPT-4 has not yet achieved the capability to generate table content with the necessary conciseness and accuracy for information summary and synthesis.

### Figure creation

In the diagram drawing test, we removed all terms indicative of a cyclical graph from the diagram description in the prompt to evaluate whether GPT-4 could independently recreate the original, pioneering depiction of the cancer immune system cycle. We employed three strategies for diagram generation, as depicted in Figure 2, which included: 1) using a diagram description generated from references and incorporated into the drawing prompt; 2) using the description from BRP2; 3) relying on the GPT-4 base model. The resulting diagrams produced by GPT-4 are presented in Figure 3, with detailed information provided in the supplementary materials.

**Figure 3.**
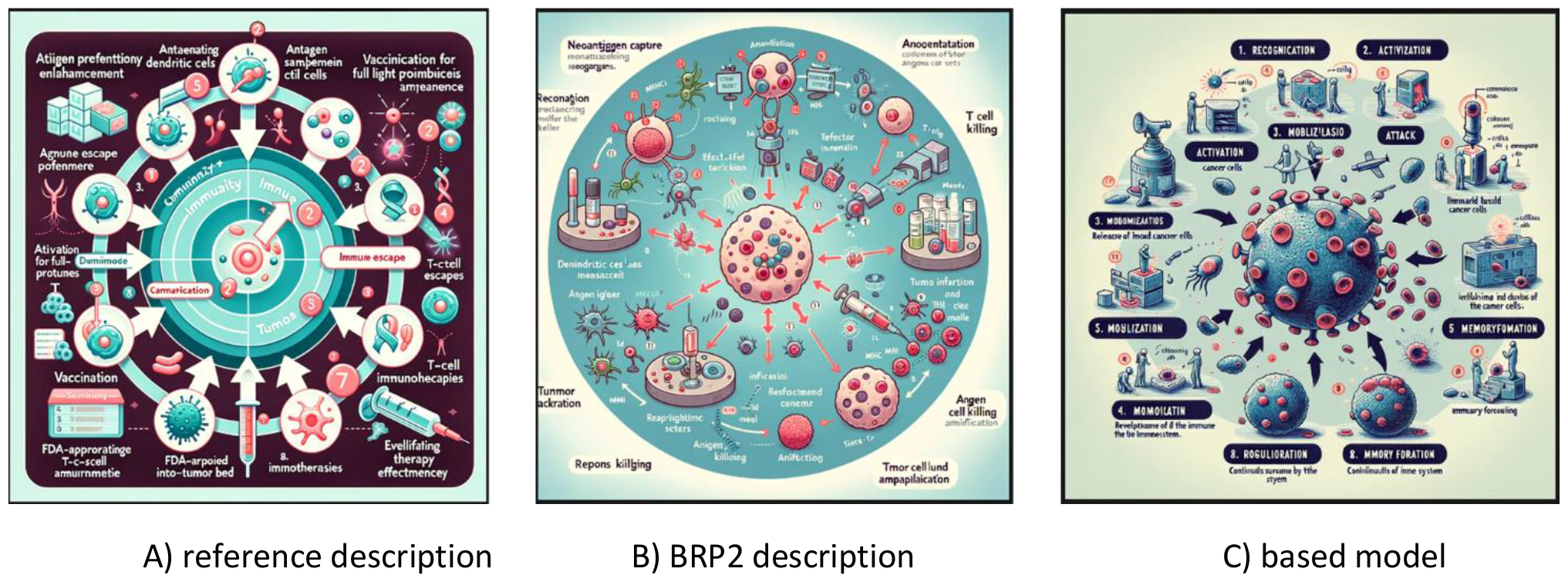

These diagrams highlight common inaccuracies in GPT-4’s drawings, such as misspelled words, omitted numbers, and a lack of visual clarity due to superfluous icons and cluttered labeling. Despite these issues, GPT-4 demonstrated remarkable proficiency in constructing an accurate cycle architecture, even without explicit instructions to do so.

In conclusion, while GPT-4 can serve as a valuable tool for conceptualizing diagrams for various biomedical reactions, mechanisms, or systems, professional graph drawing tools are essential for the actual creation of diagrams.

## Conclusions

In this study, we evaluated the capabilities of the language model GPT-4 within ChatGPT for composing a biomedical review article. We focused on four key areas: (1) summarizing insights from reference papers; (2) generating text content based on these insights; (3) suggesting avenues for future research; and (4) creating tables and graphs. GPT-4 exhibited commendable performance in the first three tasks but was unable to fulfill the fourth.

ChatGPT’s design is centered around text generation, with its language model finely tuned for this purpose through extensive training on a wide array of sources, including scientific literature.

Consequently, GPT-4’s proficiency in text summarization and synthesis is anticipated. Remarkably, the GPT-4 base model’s performance is on par with, or in some cases, slightly surpasses that of reference-based text content generation, owing to its training on a diverse collection of research articles and web text. Furthermore, reproducibility tests have demonstrated GPT-4’s ability to generate consistent text content, whether utilizing references or solely relying on its base model.

In addition, we assessed GPT-4’s proficiency in extracting precise and pertinent information for the construction of research-related tables. GPT-4 encountered difficulties with this task, indicating that ChatGPT’s language model requires additional training to enhance its ability to discern and comprehend specialized scientific terminology from literature. This improvement necessitates addressing complex scientific concepts and integrating knowledge across various disciplines.

Moreover, GPT-4’s capability to produce scientific diagrams does not meet the standards required for publication. This shortfall may stem from its associated image generation module, DALL-E, being trained on a broad spectrum of images that encompass both scientific and general content. However, with ongoing updates and targeted retraining to include a greater volume of scientific imagery, the prospect of a more sophisticated language model with improved diagrammatic capabilities could be a foreseeable advancement.

To advance the assessment of ChatGPT’s utility in publishing biomedical review articles, we executed a plagiarism analysis on the text generated by GPT-4. This analysis revealed potential issues when references were employed, with GPT-4 occasionally producing outputs that closely resemble content from reference articles. Although GPT-4 predominantly generates original text, we advise conducting a plagiarism check on ChatGPT’s output before any formal dissemination. Moreover, despite the possibility that the original review paper BRP1 was part of GPT-4’s training dataset, the plagiarism evaluation suggests that the output does not unduly prioritize it, considering the extensive data corpus used for training the language model.

Our study also highlights the robust performance of the GPT-4 base model, which shows adeptness even without specific reference articles. This observation leads to the conjecture that incorporating the entirety of scientific literature into the training of a future ChatGPT language model could facilitate the on-demand extraction of review materials. Thus, it posits the potential for ChatGPT to eventually author comprehensive summary and synthesis-based scientific review articles.

ChatGPT’s power and versatility warrant additional exploration of various facets. While these are beyond the scope of the current paper, we will highlight selected topics that are instrumental in fostering a more science oriented ChatGPT environment.

- Holistic evaluation

To thoroughly assess ChatGPT’s proficiency in generating biomedical review papers, it is imperative to include a diverse range of review paper types in the evaluation process. For instance, ChatGPT is already equipped to devise data analysis strategies and perform data science tasks in real-time. This capability suggests potential for generating review papers that include performance comparisons and benchmarks of computational tools. However, this extends beyond the scope of our pilot study, which serves as a foundational step toward more extensive research endeavors.

- Statistics of reference documents

Ideally, ChatGPT would conduct essential statistical analyses of uploaded documents, such as ranking insights, categorizing documents per insight, and assigning relevance weights to each document. This functionality would enable scientists to quickly synthesize the progression and extensively studied areas within a field.

- Mitigating hallucination

Employing uploaded documents as reference material can reduce the occurrence of generating inaccurate or ‘hallucinated’ content. However, when queries exceed the scope of these documents, ChatGPT may still integrate its intrinsic knowledge base. In such cases, verifying ChatGPT’s responses against the documents’ content is vital. A feasible method is to cross-reference responses with the documents, although this may require significant manual effort. Alternatively, requesting ChatGPT to annotate its output with corresponding references from the documents could be explored, despite being a current limitation of GPT-4.

- Academic Integrity concerns

As the development of LLMs progresses towards features that could potentially expedite or even automate the creation of scientific review papers, the establishment of a widely accepted ethical practice guide becomes paramount. Until such guidelines are in place, it remains essential to conduct plagiarism checks on AI-generated content and transparently disclose the extent of AI’s contribution to the published work.

- Comparison with other LLM models

The advent of large language models like Google’s Gemini AI [26] and Perplexity.ai has showcased NLP capabilities comparable to those of GPT-4. This, coupled with the emergence of specialized models such as BioBert [27], BioBART [28], and BioGPT [29] for biomedical applications, highlights the imperative for in-depth comparative studies. These assessments are vital for identifying the optimal AI tool for particular tasks, taking into account aspects such as multimodal functionalities, domain-specific precision, and ethical considerations. Conducting such comparative analyses will not only aid users in making informed choices but also promote the ethical and efficacious application of these sophisticated AI technologies across diverse sectors, including healthcare and education.

## Supplements

All supplementary materials are available at https://github.com/EpistasisLab/GPT4_and_Review.

